# Psychological well-being modulates neural synchrony during naturalistic fMRI

**DOI:** 10.1101/2023.09.29.560216

**Authors:** K. Klamer, J. Craig, K. Sullivan, C. Haines, C. Ekstrand

## Abstract

Psychological well-being (PWB) is a combination of self-acceptance, life purpose, personal growth, positive relationships, and autonomy, and has a significant relationship with physical and mental health. Previous studies using resting-state functional magnetic resonance imaging (fMRI) and static picture stimuli have implicated the anterior cingulate cortex (ACC), posterior cingulate cortex (PCC), orbitofrontal cortex (OFC), insula and thalamus in PWB, however, the replication of associations across studies is scarce, both in strength and direction, resulting in the absence of a model of how PWB impacts neurological processing. Naturalistic stimuli better encapsulate everyday experiences and can elicit more “true-to-life” neurological responses, and therefore may be a more appropriate tool to study PWB. The current study seeks to identify how differing levels of PWB modulate neural synchrony in response to an audiovisual film. With consideration of the inherent variability of the literature, we aim to ascertain the validity of the regions previously mentioned and their association with PWB. We identified that higher levels of PWB were associated with heightened neural synchrony in the bilateral OFC and left PCC, and that lower levels of PWB were associated with heightened neural synchrony in the right temporal parietal junction (TPJ) and left superior parietal lobule (SPL), regions related to narrative processing. Taken together, this research confirms the validity of several regions in association with PWB and suggests that varying levels of PWB produce differences in the processing of a narrative during complex audiovisual processing.

## Psychological well-being modulates neural synchrony during naturalistic fMRI

Well-being refers to optimal functioning (Lyubomirsky et al., 2005; Ryan & Deci, 2001) and is a complex trait-like construct that has evolved over time. The investigation of well-being generally asks what it means to live a good life, and the framework for understanding well-being historically distinguishes between hedonic and eudaimonic well-being (King, 2019). Eudaimonic well-being relates to the consequences of self-growth and self-actualization, whereas hedonic well-being relates to immediate sensory pleasure, happiness and enjoyment (Ryan & Deci, 2001; Ryff, 1989). In the contemporary investigation of well-being, the most frequently used terms are subjective well-being (SWB) and psychological well-being (PWB), which are thought to respectively originate from hedonic and eudaimonic traditions (King, 2019). PWB and SWB are correlated constructs, where SWB involves life satisfaction, the absence of negative emotions and the presence of positive emotions (Hausler et al., 2017), where PWB incorporates experiencing positive relationships, life purpose and the development of potential (Huppert, 2009). Although some researchers consider them to be distinct, current models suggest strong correlation and a lack of discriminant validity between the two terms (King, 2019). PWB is therefore seen as an umbrella term encompassing hedonic and eudaimonic aspects of well-being, aligning with theoretical perspectives (Salsman et al., 2014). In other words, PWB considers life satisfaction, positive/negative mental and emotional states (e.g., feelings of happiness, sadness, etc), and sense of purpose/meaning in life (Steptoe et al., 2015).

While research generally focuses on neurological mechanisms underlying psychopathology, the past 60 years has demonstrated an increase in the literature devoted to the study of well-being (King, 2019), in part due to the discovery that higher PWB is linked to better physical and mental health outcomes (Huppert, 2009). Specifically, evidence has suggested that positive hedonic states, eudaimonic well-being, and life evaluation are relevant to health and quality of life as people age (Steptoe et al., 2015). It has been well established that impaired PWB is associated with increased risk of physical illness due to the long-standing acceptance that psychological distress is strongly correlated with reduced quality and duration of life, increased use of health services, and physical morbidity (Winefield et al., 2012). However, new evidence has shown that positive PWB could be a protective factor for health (Lyubomirsky et al., 2005), as studies have suggested that positive hedonic states and life evaluations predict lower future mortality and morbidity (Chida & Steptoe, 2008). Positive PWB is associated with reduced cortisol output during the day (Steptoe et al., 2005; Steptoe et al., 2008), which may impact immune regulation and lipid metabolism (Winefield et al., 2012). Further, positive affect, which is an aspect of hedonic well-being, has been associated with reduced inflammatory and cardiovascular responses to acute mental stress, and the reduction in inflammatory markers (Steptoe et al., 2012). Moreover, PWB is associated with attaining and maintaining higher physical activity levels (Kim et al., 2017) and has a significant effect on mental health (Salsman et al., 2014). PWB can therefore impact physical health both directly and indirectly through the effects of positive or negative mental health states.

PWB has a broad influence on cognition across various domains. PWB may influence attention processes, as evidenced by a meta-analysis (Irie et al., 2019) reporting that attention is related to positive aspects of mental health, including well-being. Further, mindfulness, which is characterized by purposeful control of attention (Brown et al., 2007, is highly positively correlated with PWB and negatively correlated with psychological distress (Parto & Besharat, 2011). PWB has been shown to have links to memory processes, specifically semantic self-images, and this correlation grows stronger as age increases (Rathbone et al., 2015). Semantic self-images have been suggested to play a crucial role in supporting the self, providing a link between semantic memory and eudaimonic processes of PWB (Rathbone et al., 2015). Studies have also suggested that PWB has a positive significant correlation with adaptive decision-making strategies in adolescents, particularly with subjective aspects of PWB. Higher levels of PWB are associated with a preference for adaptive decision-making strategies, emphasizing life satisfaction and self-realization (Páez-Gallego et al., 2020). Likewise, cognitive flexibility has been found to statistically predict PWB, where it is considered a mediator in the relationship between self-confidence and PWB (Malkoç & Kesen Mutlu, 2019). Importantly, a significant relationship has been observed between PWB and aging, particularly in the health outcomes at older ages (Steptoe et al., 2015). PWB may contribute to longer life expectancies and promote active aging through a protective role (McFarquhar & Bowling, 2009), and attitudes toward aging may moderate the relationship between PWB and subjective age (Mock & Eibach, 2011). Therefore, the impact of PWB on behaviour and cognition is far reaching and may have profound effects on mental and physical health outcomes.

While an abundance of neuroimaging studies have investigated the neural correlates of PWB, there is a large degree of variability in the literature. As reviewed by King (2019), the anterior cingulate cortex (ACC), posterior cingulate cortex (PCC), orbitofrontal cortex (OFC), insula, and thalamus are the regions most consistently reported in association with PWB. However, there have been a wide range of brain regions suggested to be involved in PWB, and replication of associations across studies is scarce, both in strength and direction (de Vries et al., 2023). Further, the functionality of these brain regions in determining PWB remains unclear. As a result, there is an overall lack of understanding of how different levels of PWB impact or are impacted by specific brain regions, and there is no empirically validated model of PWB in the brain.

One source of stimuli that show promise for characterizing how specific personality traits modulate neural brain activity are naturalistic stimuli. Naturalistic paradigms use rich, multimodal dynamic stimuli that represent our daily lived experience, such as audiovisual films and virtual reality (Sonkusare et al., 2019). Consequently, naturalistic stimuli can evoke brain responses that are highly reproducible within and across subjects (Hasson et al., 2004). This allows naturalistic stimuli to engage a broader set of brain regions and more diverse modes of network interactions than task or resting-state paradigms (Zhang et al., 2021). Finn et al. (2018) used naturalistic stimuli to characterize differences in brain synchrony between participants with low trait paranoia and high trait paranoia. Their results showed characteristic patterns of neural synchrony unique to the high paranoia group, suggesting that personality traits can influence the processing of naturalistic stimuli in the brain. Based on this, naturalistic paradigms may be an effective tool for examining how PWB modulates neural activity and may help create a model of PWB in the brain.

While viewing a naturalistic audiovisual stimulus, the narrative of the story influences the neural activity in response, and higher ISC seems to predict the efficacy of the narrative. A review by Jääskeläinen et al. (2020) unravels the findings from neuroimaging studies on how narratives influence the human brain, and suggested that distinct regions in the brain have distinct roles when it comes to the processing of a narrative. ISC in the temporoparietal junction (TPJ) is associated with higher suspense, attentionally engaging narratives produce synchronous activity in the intraparietal sulcus/superior parietal lobule (IPS/SPL) and the frontal eye field (FEF), ISC in the precuneus is associated with narrative sense-making, and narratives eliciting negative emotions synchronize both the precuneus and the ventromedial prefrontal cortex (vmPFC). Lastly, visual and auditory cortices, such as the lateral occipital cortex (LOC), superior temporal gyrus (STG), show synchronous activity due to the sensory processing of an identical audiovisual stimulus. As a result, we can expect to see synchronous responses in the aforementioned brain regions during any naturalistic audiovisual movie viewing task, highlighting the intricate coordination of these regions in integrating visual and auditory cues, processing different person identities, and enabling the multimodal integration required to construct a coherent, continuous narrative.

The current study seeks to investigate differential neural synchrony associated with varying levels of PWB using audiovisual movies during fMRI. We used preprocessed data from the Naturalistic Neuroimaging Database (Aliko et al., 2020, v2.0) from 20 participants who viewed a feature length audiovisual movie during fMRI. PWB for each participant was quantified using the NIH Toolbox 2.0 Psychological Well-Being Instrument (Gershon et al., 2013). We hypothesize that, in line with previous research, varying levels of PWB will be associated with differential neural synchrony in the OFC, ACC, PCC, thalamus and insula. We additionally hypothesize there will be differential neural synchrony in regions associated with narrative processing, including the SPL/IPS, FEF, TPJ, LOC and STG. With consideration of the inherent variability in the extant literature on PWB, this study is framed as an exploratory inquiry, and we aim to find consistent neural synchrony associated with low and high levels of PWB. By using naturalistic stimuli, we anticipate that we may uncover novel differences in how differing levels of PWB impact complex multimodal integration.

## Methods

### Participants and fMRI data

We used fMRI data from the publicly available Naturalistic Neuroimaging Database (NNDb, v2.0; Aliko et al., 2020). We selected 20 participants (10 females/10 males, aged 19-53, mean age of 27.7 years) who watched the feature-length audiovisual film *500 Days of Summer* (Webb, 2009; duration ∼ 95 minutes) during fMRI. All participants were right-handed, native English speakers, with no history of neurological/psychiatric illnesses, no hearing impairments, unimpaired or corrected vision, and did not take medication. Full fMRI preprocessing details can be found in Aliko et al. (2020). Briefly, all data was acquired on a 1.5T Siemens MAGNETOM Avanto, and functional data was acquired using a multiband echo-planar imaging (EPI) sequence (TR: 1s, TE: 58.4ms). We used version 2.0 of the NNDb (Aliko et al., 2020), which, in comparison to v1.0, has improved normalization and standardization of the data, resulting in ‘cleaner’ preprocessed data and more robust statistics. Aliko et al., (2020) used the *afni_proc.py* pipeline from AFNI (Cox & Hyde, 1997; Cox, 1999) for preprocessing, correcting for slice-timing differences, despiking, correcting for motion, spatially aligning the data to an MNI template with a resampling size of 3×3×3mm^3^, and correcting for timing to align the fMRI time series and the film. The authors of the NNDb (v2.0) (Aliko et al., 2020) obtained approval by the ethics committee of University College London and participants provided written informed consent to take part in the study and share their anonymized data.

### Behavioural Questionnaires

Following the MRI imaging session, participants completed the majority of the National Institute of Health (NIH) Toolbox, which validates measures of sensory, motor, cognitive and emotional processing to measure individual differences (Gershon et al., 2013). In this study, we used the Psychological Well-Being (PWB) scores quantified using the NIH Toolbox 2.0 Psychological Well-Being (18+; Gershon et al., 2013) questionnaire. We used a median split analysis to separate participants into low PWB and high PWB groups based on their provided T-score, resulting in 10 participants in each group. We used SPSS (version 27; IBM Corp., 2020) to run an independent sample t-test between participants in each PWB group (i.e., low-low and high-high PWB) and age.

### Data Availability Statement

The preprocessed fMRI data used in this study is openly available and can be downloaded from the Naturalistic Neuroimaging Database (version 2.0.0; https://openneuro.org/datasets/ds002837/versions/2.0.0).

### Intersubject Correlation Analyses

Intersubject correlation analyses (ISC) were run for all unique pairs of participants, which produced 190 (*n*\*(*n*-1)/2, where n = 20) unique ISC maps. ISC is a model-free approach used to analyze complex fMRI data acquired in naturalistic, audiovisual stimulus environments (Kauppi et al., 2010). It allows us to measure shared content across experimental conditions by filtering out subject-specific signals and revealing voxels with a consistent, stimulus-evoked response time series across subjects (Nastase et al., 2019). It does this by calculating pairwise correlation coefficients between all pairs of participants for each voxel throughout the brain. We separated the resulting 190 ISC maps into three groups based on PWB grouping (i.e., low-low PWB pairwise correlations grouped, high-high pairwise correlations grouped, low-high pairwise correlations grouped). This resulted in 45 (*n*\*(*n*-1)/2, where n = 10) unique ISC maps for both the high (i.e., high-high) and low (i.e., low-low) PWB groups, and 100 between group pairs (i.e., low-high).

### FMRI Analysis

After running ISCs on all pairs of participants, we used the Linear Mixed Effects Model (LME) implemented via the *3dISC* module in AFNI (Cox & Hyde, 1997; Cox, 1999; as described by Chen et al., 2020). LME is a parametric method based on the general linear model (GLM) and is used to model the relationship between the fMRI time series data and the experimental conditions at each voxel, accounting for the complex covariance structure of ISC data. As LME accounts for variability in the data due to random effects, it allows for more precise estimation of fixed effects (Koerner & Zhang, 2017). We modeled the following contrasts: ISC in the low PWB group (low-low PWB), ISC in the high PWB group (high-high PWB), ISC between the high and low groups (low-high PWB), comparison of ISC between the low and high groups (low-low vs. high-high PWB), comparison of ISC between the low group and between group ISC (low-low vs. low-high), and comparison of ISC between the high group and between group ISC (high-high vs. low-high). Significant ISCs for the group average contrasts (i.e.., high-high PWB and low-low PWB), were defined using a voxelwise false discovery rate (FDR) of *q* < 0.0001. Significant ISCs for the group comparisons contrasts (i.e., low-low PWB vs. high-high PWB; low-low PWB vs. low-high PWB; high-high vs. low-high PWB) were defined using a voxelwise FDR of *q* < 0.05. Significant results were transformed into surface space for visualization purposes only.

## Results

### Behavioural Questionnaires

We used SPSS (version 27; IBM Corp., 2020) to run an independent sample t-test between participants in the low and high PWB groups to determine whether age differed between the two groups. No significant differences were found *t*(18) = .388, *p* = .703.

### Group Averages

Results from the low-low PWB and high-high PWB group average contrasts showed widespread ISCs across occipital, temporal, frontal, and parietal cortices, which is in line with previous research (Güçlütürk et al., 2018). Full results for low-low PWB and high-high PWB group averages can be found in Supplementary Materials A.

### Low-low vs. High-high PWB contrasts

#### Low-low > High-high PWB

Results from the low-low PWB > high-high PWB are shown in Figure 1 and Supplementary Materials B. The Low-low PWB group showed significantly greater synchrony than the High-high PWB group in the bilateral LOC, bilateral supracalcarine cortex, bilateral insula, bilateral precuneus, right IPS, right TPJ, and right frontal eye field (FEF).

**Figure 1.**
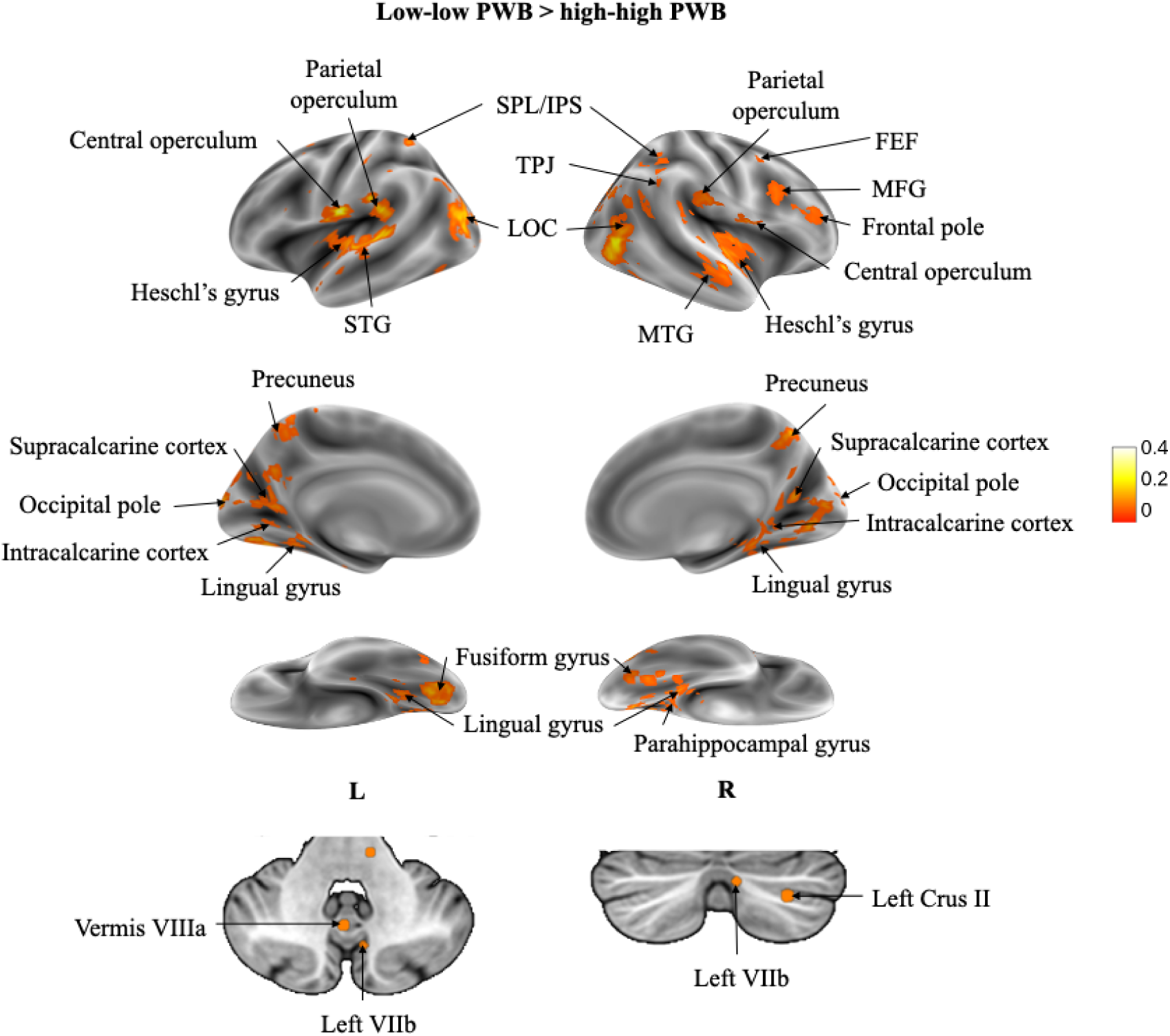
Voxels showing significant ISC within participants in the low-low PWB compared to the high-high PWB group across the time course of the audiovisual stimulus. Results are displayed as a voxelwise false-discovery rate (FDR) threshold of *q* < 0.05.

#### High-high > Low-low PWB

Results from the High-high > Low-low PWB contrast are shown in Figure 2 and Supplementary Materials B. Overall, the high-high PWB group showed significantly greater synchrony than the low-low PWB group in the right SPL, bilateral OFC, bilateral PCC, and right precentral gyrus.

**Figure 2.**
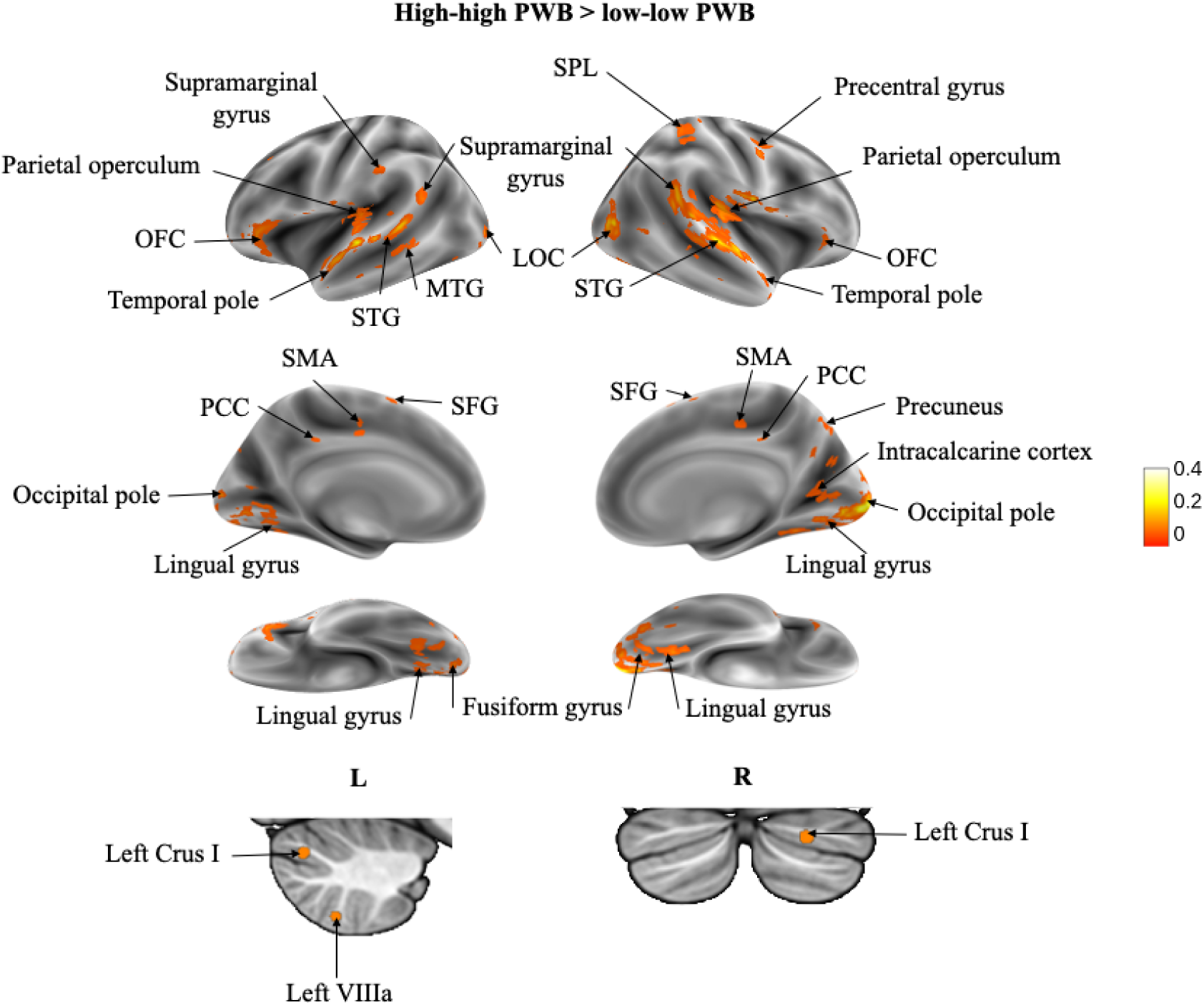
Voxels showing significant ISC within participants in the high-high PWB compared to the low-low PWB group across the time course of the audiovisual stimulus. Results are displayed as a voxelwise false-discovery rate (FDR) threshold of *q* < 0.05.

### Comparing low and high group ISC with between group ISC

#### Low-low > Low-high PWB

Results from the Low-low > Low-high PWB contrast are shown in Figure 3 and Supplementary Materials C. This contrast showed heightened neural synchrony in comparison to the low-high PWB group in the left SPL, right TPJ, right middle frontal gyrus (MFG), left parietal operculum, and the right frontal pole.

**Figure 3.**
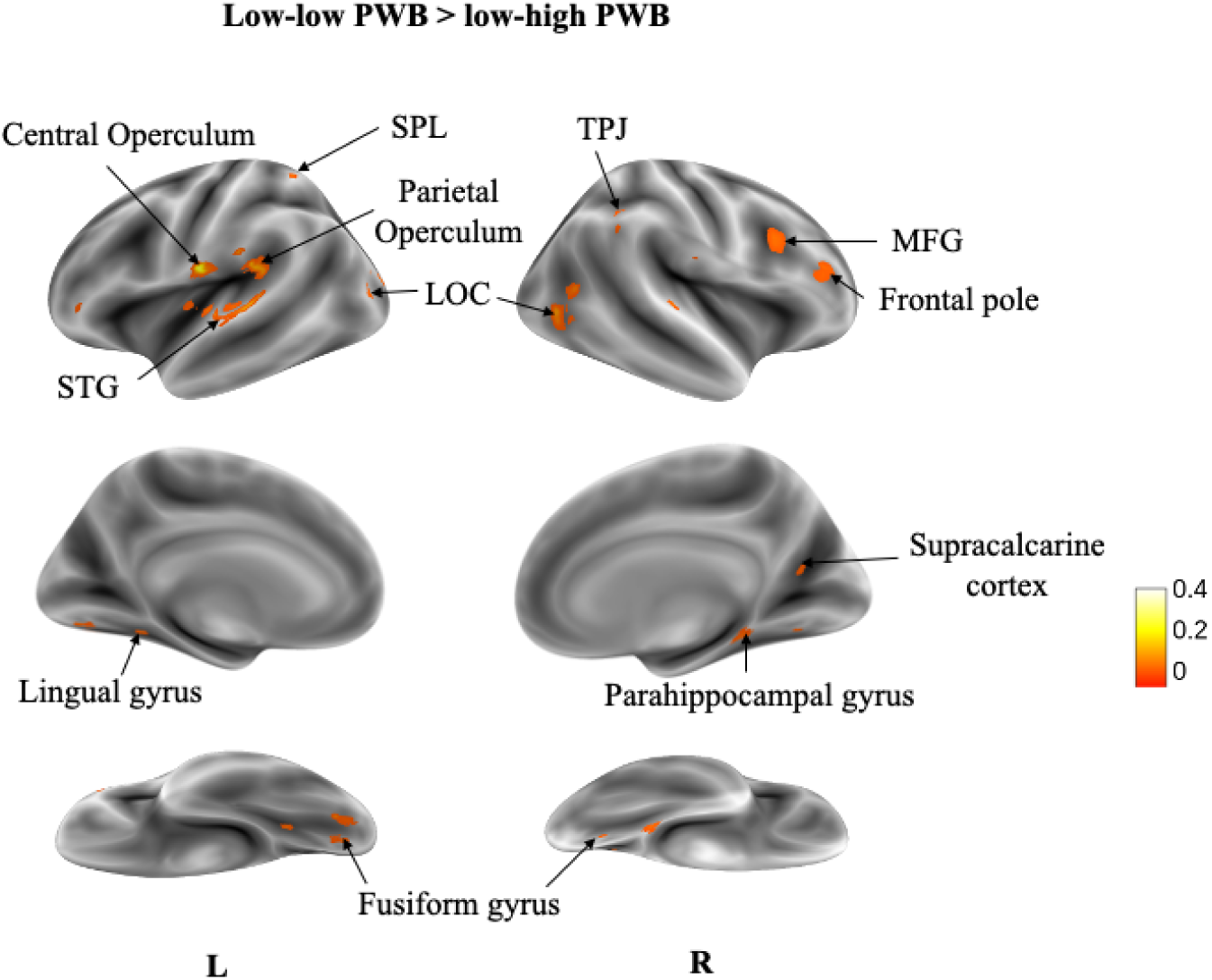
Voxels showing significant ISC across the time course of the audiovisual stimulus in participants within the low-low PWB group in comparison to participants within the low-high PWB group. Results are displayed as a voxelwise false-discovery rate (FDR) threshold of *q* = 0.05.

#### High-high > Low-high PWB

Results from the High-high > Low-high PWB contrast are shown in Figure 4 and Supplementary Materials C. Overall, the high-high PWB group showed heightened neural synchrony in comparison to the low-high PWB group in the bilateral OFC, left PCC, right STG, bilateral superior frontal gyrus (SFG), right parietal operculum, and right precentral gyrus.

**Figure 4.**
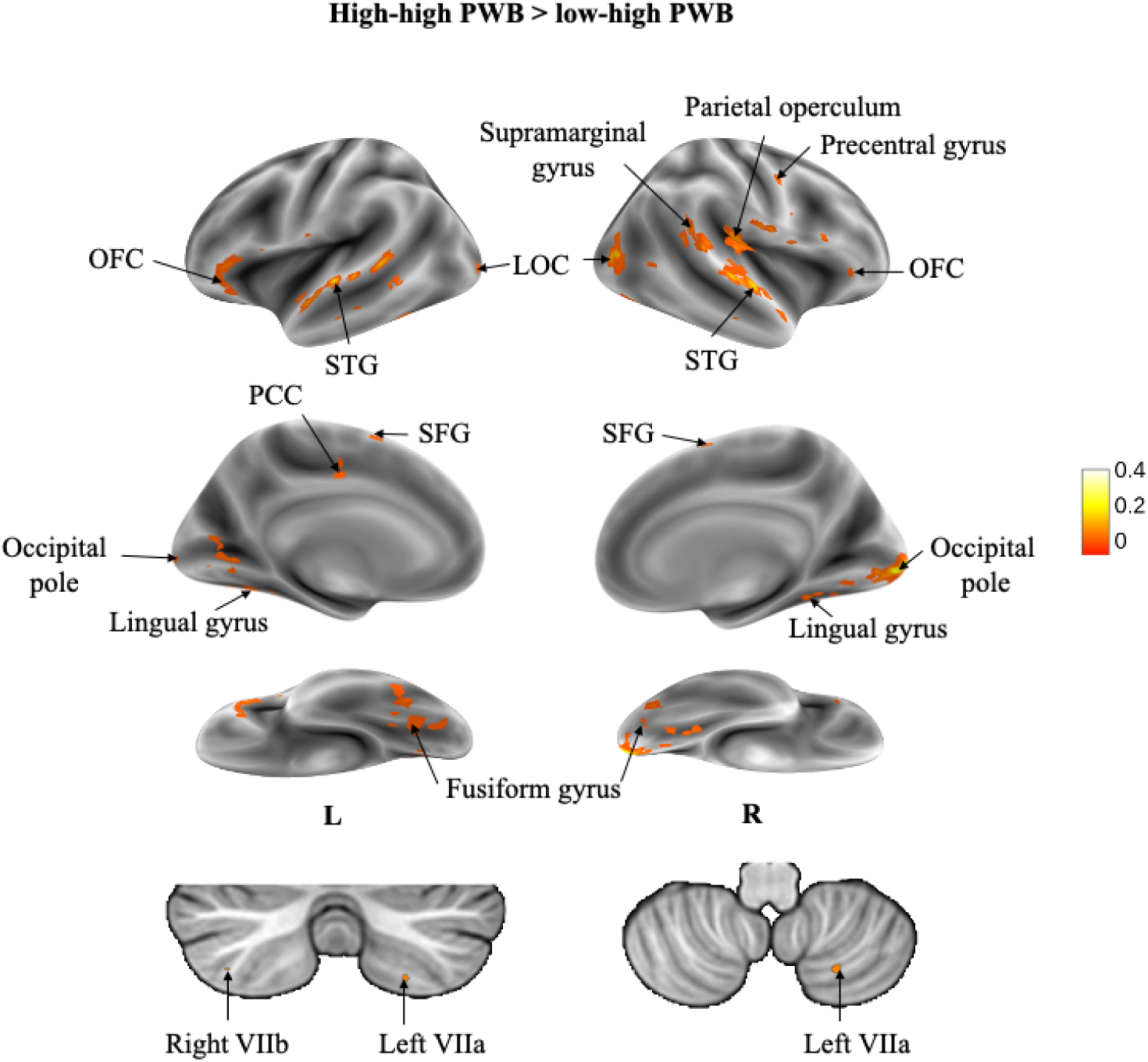
Voxels showing significant ISC across the time course of the audiovisual stimulus in participants within the high-high PWB group in comparison to participants within the low-high PWB group. Results are displayed as a voxelwise false-discovery rate (FDR) threshold of *q* = 0.05.

This study sought to examine differences in neural synchrony to a complex audiovisual stimulus based on differing levels of PWB. To do so, we separated participants into low or high PWB groups using a median split analysis, ran ISC analysis and used LME modelling to visualize how neural synchrony differs between each group. Neural activity has been shown to be sensitive to the nature of the stimulus, and results from this study have confirmed that individual differences in PWB result in substantial variations in neural synchrony in response to a naturalistic stimulus. Consistent with our hypothesis, differential neural synchrony was found in association with varying levels of PWB in regions associated with PWB and narrative processing.

The left TPJ and right SPL were found to have higher synchronization in the high-high PWB group compared to the low-low and low-high PWB groups. During narrative processing, the TPJ is centrally associated with higher synchronization during suspenseful narratives and negative emotional narratives (Jääskeläinen et al., 2020). Therefore, higher synchronization in this region may indicate that individuals with lower levels of PWB found the narrative more suspenseful and interpreted it more negatively. Previous research has demonstrated that psychological distress, which is in part defined by low PWB, is associated with perceiving stressful events as a negative turning point (Sutin et al., 2010). If we extend this finding to our result, this could indicate that individuals with lower levels of PWB interpreted stressful events during the film as a negative turning point, causing the more suspenseful and negative interpretation of the narrative. The SPL has been shown to have higher synchronization during narrative processing during attentionally engaging narratives (Jääskeläinen et al., 2020). This may indicate that individuals in the low-low PWB group found that narrative more attentionally engaging than individuals with higher levels of PWB. Previous research has suggested that low PWB is associated with higher attentional engagement towards emotional stimuli through its associations with low self-esteem. Low self-esteem is related to several psychiatric disorders (Shen, 2007; Silverstone & Salsali, 2003) and is therefore associated with low PWB outcomes, and there is an attention bias for negative stimuli among individuals with low self-esteem (Li et al., 2011) with the greater mobilization of attentional resources toward emotional stimuli (Li & Yang, 2013). Taken together, these results suggest that individuals with lower levels of PWB may have assigned a more negative interpretation to stressful events during the film, and therefore exhibited greater attentional engagement throughout the film.

The right MFG and right frontal pole also showed higher synchronization in the low-low PWB group compared to the low-high and high-high PWB groups. These results indicate that these regions responded more similarly in the low-low PWB group, suggesting that these regions play a role in mediating lower PWB outcomes in response to an audiovisual stimulus. These regions have not been previously associated with PWB; therefore, more research is needed to fully elucidate the role of the right MFG and right frontal pole in PWB.

The regions that showed higher synchronization in the high-high PWB group compared to the low-low and low-high PWB groups include the bilateral OFC and left PCC. The OFC is a region that has been previously associated with PWB, with a resting-state fMRI study finding that the remote cortical connectivity of the right OFC was positively correlated with PWB (Li et al., 2022), and grey matter volume of the left OFC being associated with increased optimism (i.e., a component of positive affect; Dolcos et al., 2016). Further, the OFC is a key brain area in the representation of reward value (Rolls et al., 2020), providing a direct link to subjective components of PWB. Higher synchronization of this region suggests that it behaved more similarly in participants with higher PWB throughout the audiovisual film, emphasizing previous reports that associate the OFC with higher PWB outcomes. The PCC has previously been consistently associated with PWB (King, 2019), however, the nature of how it mediates differential PWB outcomes has not yet been determined. One plausible theory is that the PCC may mediate mindfulness, which is associated with higher PWB outcomes. Previous research has indicated that the PCC shows significant effects related to meditation and mindfulness (Zsadanyi et al., 2021) and has implicated the PCC as a plausible mechanistic target for meditation (Brewer & Garrison, 2014). The finding in the current study of heightened synchrony in the PCC suggests that the PCC behaved more similarly in individuals with higher levels of PWB in response to the audiovisual film and provides support for previous research associating the PCC and higher PWB outcomes. While it is plausible that this heightened synchrony was mediated through increased mindfulness during the film, more research is needed to fully determine the role of the PCC in mediating higher PWB outcomes.

Heightened synchrony was consistently found in the right precentral gyrus in the high-high PWB group compared to the low-low and low-high PWB groups. This indicates that the right precentral gyrus responded more similarly throughout the film in individuals with higher levels of psychological well-being, suggesting an association between this region and higher PWB outcomes. However, this region has not been previously associated with differential PWB, therefore, more research is needed to elucidate the relationship between higher levels of PWB and the right precentral gyrus.

This study aimed at uncovering differential neural synchrony associated with varying levels of PWB levels during movie watching, however, there are several limitations to be discussed. Due to the use of an online database, we were unable to have control over the stimuli that were presented to participants. This introduces a limitation as previous literature that suggests the valence of the stimulus may alter the neural response based on levels of PWB. As the database utilized has only two films with a sample size larger than *n* = 6, one being a documentary, we were limited to only one plausible choice when determining which film would be used as our stimulus. Additionally, the data used in these experiments was collected using a 1.5T MRI, which has a lower signal-to-noise ratio and a lower spatial specificity than MRIs with a stronger field strength. Future research should aim at using several audiovisual films of differing emotional valence and should be investigated using higher field strengths to acquire such data.

In conclusion, this study demonstrated that variations in levels of PWB may modulate differential neural synchrony during movie-watching. We confirmed previous research that implicated several regions in higher levels of PWB, including the bilateral OFC and left PCC, regions that relate to cognitive traits such as mindfulness and the determination of subjective reward value. Additionally, we found novel differences associated with lower levels of PWB in the left TPJ and right SPL, regions related to various aspects of narrative processing. Due to the use of a naturalistic paradigm, these results can be applied to real-life cognition, and collectively emphasize the profound effect that differing levels of PWB have on complex, multimodal processing.

