## Supplementary Materials for "Psychological well-being modulates neural synchrony during naturalistic fMRI"

**Supplementary Materials A**

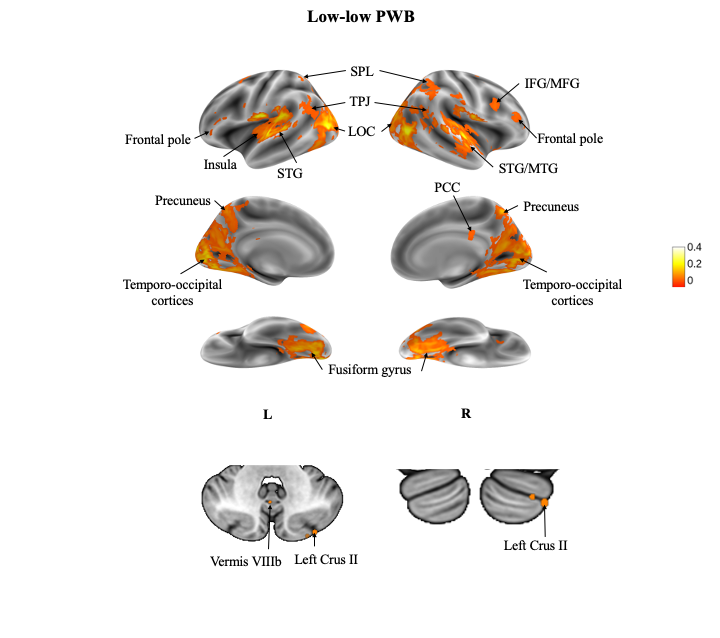

**Figure 1.** Voxels showing significant ISC across the time course of the audiovisual stimulus in participants within the low-low PWB group (*n* = 10). Results are displayed as a voxelwise false-discovery rate (FDR) threshold of *q* = 0.0001.

**Table 1**

*Coordinates and cluster sizes of neural synchrony found within the low-low PWB group average contrast. Clusters with five or more voxels shown.*

| **Anatomical Location** | **Hemisphere** | **# of Voxels** | **MAX *r*** | **MAX x** | **MAX y** | **MAX z** |
| --- | --- | --- | --- | --- | --- | --- |
| Intracalcarine Cortex | Left | 941 | 0.242 | -10.5 | -82.5 | 7.5 |
| Lateral Occipital Cortex, inferior division | Right | 330 | 0.256 | 43.5 | -64.5 | 4.5 |
| Lateral Occipital Cortex, inferior division | Left | 230 | 0.296 | -52.5 | -73.5 | 7.5 |
| Central Opercular Cortex | Left | 228 | 0.407 | -61.5 | -16.5 | 10.5 |
| Occipital Fusiform Gyrus | Right | 184 | 0.138 | 22.5 | -73.5 | -10.5 |
| Planum Temporale | Right | 177 | 0.294 | 58.5 | -10.5 | 4.5 |
| Precuneous Cortex | Right | 80 | 0.147 | 1.5 | -58.5 | 52.5 |
| Lateral Occipital Cortex, superior division | Right | 67 | 0.0838 | 34.5 | -58.5 | 58.5 |
| Lateral Occipital Cortex, inferior division | Left | 31 | 0.114 | -43.5 | -67.5 | 1.5 |
| Angular Gyrus | Left | 27 | 0.0904 | -55.5 | -61.5 | 22.5 |
| Supramarginal Gyrus, posterior division | Right | 26 | 0.0902 | 46.5 | -43.5 | 19.5 |
| Lateral Occipital Cortex, superior division | Right | 23 | 0.079 | 25.5 | -73.5 | 49.5 |
| Inferior Frontal Gyrus, pars opercularis | Right | 11 | 0.0553 | 37.5 | 10.5 | 25.5 |
| Superior Temporal Gyrus, posterior division | Right | 9 | 0.18 | 64.5 | -28.5 | 1.5 |
| Precuneous Cortex | Right | 7 | 0.0324 | 1.5 | -55.5 | 22.5 |
| Lateral Occipital Cortex, superior division | Right | 7 | 0.102 | 10.5 | -61.5 | 67.5 |
| Middle Frontal Gyrus | Right | 7 | 0.0546 | 43.5 | 34.5 | 19.5 |
| Inferior Temporal Gyrus, temporooccipital part | Right | 6 | 0.0835 | 49.5 | -58.5 | -10.5 |
| Lateral Occipital Cortex, superior division | Left | 6 | 0.101 | -16.5 | -79.5 | 46.5 |
| Angular Gyrus | Right | 6 | 0.0813 | 58.5 | -49.5 | 13.5 |
| Lateral Occipital Cortex, superior division | Right | 5 | 0.0676 | 52.5 | -61.5 | 22.5 |
| Lateral Occipital Cortex, superior division | Left | 5 | 0.07 | -28.5 | -70.5 | 46.5 |

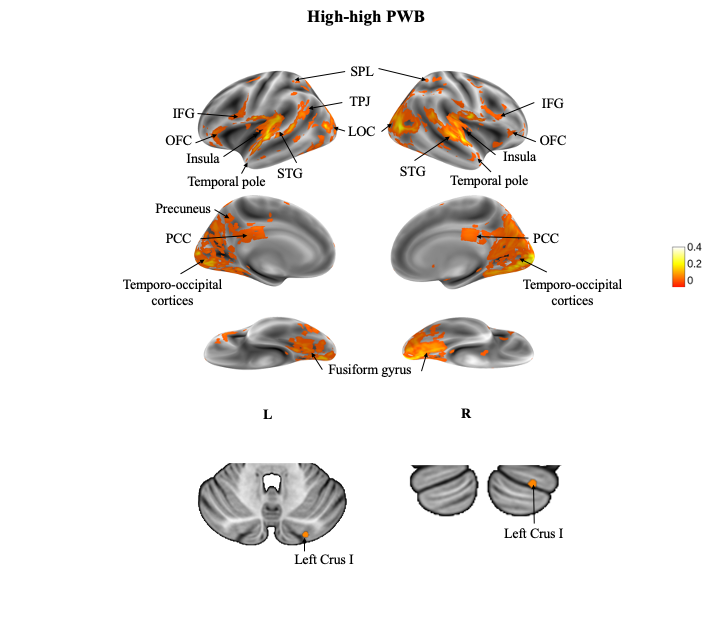

**Figure 2.** Voxels showing significant ISC across the time course of the audiovisual stimulus in participants within the low-low PWB group (*n* = 10). Results are displayed as a voxelwise false-discovery rate (FDR) threshold of *q* = 0.0001.

**Table 2**

*Coordinates and cluster sizes of neural synchrony found within the high-high PWB group average contrast. Clusters with five or more voxels shown.*

| **Anatomical Location** | **Hemisphere** | **# of Voxels** | **MAX *r*** | **MAX x** | **MAX y** | **MAX z** |
| --- | --- | --- | --- | --- | --- | --- |
| Occipital Pole | Right | 947 | 0.31 | 10.5 | -91.5 | 1.5 |
| Intracalcarine Cortex | Right | 387 | 0.131 | 13.5 | -73.5 | 13.5 |
| Superior Temporal Gyrus, anterior division | Right | 194 | 0.334 | 64.5 | -4.5 | 4.5 |
| Planum Temporale | Left | 153 | 0.349 | -64.5 | -13.5 | 4.5 |
| Lateral Occipital Cortex, inferior division | Left | 75 | 0.214 | -52.5 | -73.5 | 10.5 |
| Temporal Occipital Fusiform Cortex | Left | 72 | 0.116 | -28.5 | -55.5 | -7.5 |
| Inferior Frontal Gyrus, pars triangularis | Left | 60 | 0.0871 | -52.5 | 25.5 | -4.5 |
| Superior Temporal Gyrus, posterior division | Right | 44 | 0.155 | 70.5 | -34.5 | 10.5 |
| Superior Temporal Gyrus, posterior division | Left | 44 | 0.199 | -58.5 | -37.5 | 4.5 |
| Superior Parietal Lobule | Left | 36 | 0.0865 | -13.5 | -58.5 | 64.5 |
| Cingulate Gyrus, posterior division | Left | 25 | 0.0472 | -1.5 | -28.5 | 28.5 |
| Lateral Occipital Cortex, superior division | Right | 22 | 0.0878 | 10.5 | -64.5 | 64.5 |
| Precentral Gyrus | Left | 20 | 0.0454 | -52.5 | 7.5 | 19.5 |
| Precentral Gyrus | Right | 18 | 0.0552 | 46.5 | 10.5 | 25.5 |
| Occipital Pole | Left | 13 | 0.132 | -10.5 | -91.5 | 25.5 |
| Cingulate Gyrus, posterior division | Left | 12 | 0.0365 | -4.5 | -37.5 | 25.5 |
| Precuneous Cortex | Right | 11 | 0.0894 | 1.5 | -67.5 | 55.5 |
| Lateral Occipital Cortex, superior division | Right | 9 | 0.0463 | 43.5 | -73.5 | 43.5 |
| Occipital Fusiform Gyrus | Left | 9 | 0.0906 | -31.5 | -79.5 | -10.5 |
| Planum Polare | Left | 8 | 0.079 | -49.5 | 1.5 | -4.5 |
| Precuneous Cortex | Left | 7 | 0.0603 | -1.5 | -55.5 | 43.5 |
| Superior Parietal Lobule | Right | 7 | 0.0562 | 37.5 | -55.5 | 61.5 |
| Cingulate Gyrus, posterior division | Left | 7 | 0.0299 | -4.5 | -49.5 | 28.5 |
| Supramarginal Gyrus, posterior division | Left | 7 | 0.0318 | -55.5 | -43.5 | 49.5 |
| Occipital Pole | Right | 6 | 0.111 | 25.5 | -97.5 | -7.5 |
| Angular Gyrus | Right | 6 | 0.0349 | 43.5 | -49.5 | 46.5 |
| Lateral Occipital Cortex, superior division | Right | 5 | 0.0662 | 28.5 | -67.5 | 43.5 |
| Postcentral Gyrus | Right | 5 | 0.0485 | 64.5 | -16.5 | 28.5 |
| Lateral Occipital Cortex, superior division | Left | 5 | 0.0586 | -58.5 | -61.5 | 25.5 |
| Middle Temporal Gyrus, posterior division | Left | 5 | 0.162 | -67.5 | -25.5 | -1.5 |

**Supplementary Materials B**

**Table 3**

*Coordinates and cluster sizes of neural synchrony found within the low-low PWB group compared to the high-high PWB group. Clusters with five or more voxels shown.*

| **Anatomical Location** | **Hemisphere** | **# of Voxels** | **MAX *r*** | **MAX x** | **MAX y** | **MAX z** |
| --- | --- | --- | --- | --- | --- | --- |
| Central Opercular Cortex | Left | 104 | 0.407 | -61.5 | -16.5 | 10.5 |
| Lateral Occipital Cortex, inferior division | Left | 82 | 0.232 | -49.5 | -79.5 | 13.5 |
| Lateral Occipital Cortex, inferior division | Right | 64 | 0.256 | 43.5 | -64.5 | 4.5 |
| Central Opercular Cortex | Right | 60 | 0.212 | 55.5 | -4.5 | 4.5 |
| Lingual Gyrus | Left | 48 | 0.176 | -16.5 | -76.5 | -13.5 |
| Parahippocampal Gyrus, posterior division | Right | 24 | 0.109 | 22.5 | -37.5 | -13.5 |
| Supracalcarine Cortex | Left | 23 | 0.0888 | -22.5 | -61.5 | 16.5 |
| Intracalcarine Cortex | Right | 22 | 0.17 | 10.5 | -76.5 | 7.5 |
| Superior Parietal Lobule | Left | 21 | 0.096 | -34.5 | -52.5 | 67.5 |
| Angular Gyrus | Right | 21 | 0.0793 | 46.5 | -46.5 | 25.5 |
| Planum Temporale | Left | 17 | 0.22 | -43.5 | -37.5 | 16.5 |
| Precuneous Cortex | Right | 15 | 0.147 | 1.5 | -58.5 | 52.5 |
| Lateral Occipital Cortex, superior division | Right | 14 | 0.197 | 22.5 | -88.5 | 22.5 |
| Planum Temporale | Right | 12 | 0.204 | 61.5 | -25.5 | 16.5 |
| Inferior Frontal Gyrus, pars opercularis | Right | 11 | 0.0553 | 37.5 | 10.5 | 25.5 |
| Occipital Pole | Left | 10 | 0.218 | -4.5 | -94.5 | 16.5 |
| Precuneous Cortex | Right | 10 | 0.131 | 22.5 | -55.5 | 19.5 |
| Occipital Fusiform Gyrus | Right | 10 | 0.0853 | 22.5 | -64.5 | -13.5 |
| Lingual Gyrus | Left | 9 | 0.0907 | -19.5 | -43.5 | -10.5 |
| Lateral Occipital Cortex, superior division | Right | 9 | 0.0856 | 43.5 | -76.5 | 31.5 |
| Occipital Pole | Left | 8 | 0.158 | -34.5 | -91.5 | 10.5 |
| Lateral Occipital Cortex, superior division | Right | 8 | 0.142 | 31.5 | -79.5 | 28.5 |
| Lateral Occipital Cortex, superior division | Left | 7 | 0.122 | -10.5 | -85.5 | 43.5 |
| Lingual Gyrus | Left | 7 | 0.146 | -28.5 | -52.5 | -4.5 |
| Precuneous Cortex | Left | 7 | 0.0839 | -4.5 | -61.5 | 28.5 |
| Superior Temporal Gyrus, posterior division | Left | 7 | 0.134 | -61.5 | -19.5 | -1.5 |
| Precuneous Cortex | Left | 5 | 0.0737 | -4.5 | -49.5 | 64.5 |
| Lateral Occipital Cortex, superior division | Left | 5 | 0.07 | -28.5 | -70.5 | 46.5 |
| Intracalcarine Cortex | Right | 5 | 0.101 | 10.5 | -64.5 | 10.5 |
| Middle Frontal Gyrus | Right | 5 | 0.0546 | 43.5 | 34.5 | 19.5 |

**Table 4**

*Coordinates and cluster sizes of neural synchrony found within the high-high PWB group compared to the low-low PWB group. Clusters with five or more voxels shown.*

| **Anatomical Location** | **Hemisphere** | **# of Voxels** | **MAX *r*** | **MAX x** | **MAX y** | **MAX z** |
| --- | --- | --- | --- | --- | --- | --- |
| Occipital Pole | Right | 121 | 0.31 | 10.5 | -91.5 | 1.5 |
| Superior Temporal Gyrus, anterior division | Right | 71 | 0.334 | 64.5 | -4.5 | 4.5 |
| Inferior Frontal Gyrus, pars triangularis | Left | 65 | 0.0871 | -52.5 | 25.5 | -4.5 |
| Lateral Occipital Cortex, inferior division | Right | 49 | 0.269 | 52.5 | -70.5 | 10.5 |
| Angular Gyrus | Right | 37 | 0.146 | 61.5 | -43.5 | 16.5 |
| Superior Temporal Gyrus, anterior division | Left | 27 | 0.234 | -64.5 | -4.5 | -1.5 |
| Superior Temporal Gyrus, posterior division | Right | 21 | 0.158 | 67.5 | -34.5 | 10.5 |
| Planum Temporale | Right | 20 | 0.136 | 40.5 | -28.5 | 16.5 |
| Superior Temporal Gyrus, posterior division | Left | 13 | 0.199 | -58.5 | -37.5 | 4.5 |
| Precentral Gyrus | Right | 10 | 0.104 | 58.5 | -1.5 | 46.5 |
| Lingual Gyrus | Right | 10 | 0.119 | 31.5 | -52.5 | -4.5 |
| Planum Temporale | Left | 9 | 0.349 | -64.5 | -13.5 | 4.5 |
| Superior Temporal Gyrus, posterior division | Left | 8 | 0.169 | -67.5 | -22.5 | -1.5 |
| Occipital Fusiform Gyrus | Left | 8 | 0.0761 | -31.5 | -64.5 | -13.5 |
| Intracalcarine Cortex | Left | 7 | 0.0796 | -13.5 | -67.5 | 1.5 |
| Occipital Pole | Left | 7 | 0.102 | -16.5 | -97.5 | -1.5 |
| Occipital Fusiform Gyrus | Right | 6 | 0.0943 | 31.5 | -76.5 | -7.5 |
| Occipital Fusiform Gyrus | Left | 6 | 0.0906 | -13.5 | -88.5 | -10.5 |
| Superior Temporal Gyrus, anterior division | Right | 6 | 0.214 | 61.5 | 4.5 | -1.5 |
| Occipital Fusiform Gyrus | Left | 5 | 0.0865 | -31.5 | -76.5 | -10.5 |

**Supplementary Materials C**

**Table 5**

*Coordinates and cluster sizes of neural synchrony found within the low-low PWB group compared to the low-high PWB group. Clusters with five or more voxels shown.*

| **Anatomical Location** | **Hemisphere** | **# of Voxels** | **MAX *r*** | **MAX x** | **MAX y** | **MAX z** |
| --- | --- | --- | --- | --- | --- | --- |
| Central Opercular Cortex | Left | 28 | 0.407 | -61.5 | -16.5 | 10.5 |
| Occipital Fusiform Gyrus | Left | 5 | 0.137 | -16.5 | -79.5 | -16.5 |
| Middle Frontal Gyrus | Right | 5 | 0.0512 | 34.5 | 10.5 | 28.5 |

**Table 6**

*Coordinates and cluster sizes of neural synchrony found within the high-high PWB group compared to the low-high PWB group. Clusters with five or more voxels shown.*

| **Anatomical Location** | **Hemisphere** | **# of Voxels** | **MAX *r*** | **MAX x** | **MAX y** | **MAX z** |
| --- | --- | --- | --- | --- | --- | --- |
| Occipital Pole | Right | 59 | 0.31 | 10.5 | -91.5 | 1.5 |
| Inferior Frontal Gyrus, pars triangularis | Left | 44 | 0.0871 | -52.5 | 25.5 | -4.5 |
| Lateral Occipital Cortex, inferior division | Right | 42 | 0.269 | 52.5 | -70.5 | 10.5 |
| Superior Temporal Gyrus, anterior division | Right | 41 | 0.334 | 64.5 | -4.5 | 4.5 |
| Planum Temporale | Right | 15 | 0.136 | 40.5 | -28.5 | 16.5 |
| Angular Gyrus | Right | 12 | 0.142 | 64.5 | -46.5 | 19.5 |
| Superior Temporal Gyrus, anterior division | Left | 12 | 0.234 | -64.5 | -4.5 | -1.5 |
| Intracalcarine Cortex | Left | 8 | 0.0796 | -13.5 | -67.5 | 1.5 |
| Inferior Frontal Gyrus, pars opercularis | Right | 7 | 0.0492 | 55.5 | 16.5 | 31.5 |
| Superior Temporal Gyrus, posterior division | Left | 6 | 0.199 | -58.5 | -37.5 | 4.5 |
| Superior Temporal Gyrus, posterior division | Right | 5 | 0.158 | 67.5 | -34.5 | 10.5 |
| Planum Temporale | Left | 5 | 0.349 | -64.5 | -13.5 | 4.5 |
| Middle Temporal Gyrus, posterior division | Left | 5 | 0.162 | -67.5 | -25.5 | -1.5 |
| Occipital Fusiform Gyrus | Left | 5 | 0.0865 | -31.5 | -76.5 | -10.5 |
